# Characterization of central carbon metabolism of *Acinetobacter* sp. Tol 5 based on ^13^C-metabolic flux analysis

**DOI:** 10.64898/2026.09.29.755562

**Authors:** Erina Miyamoto, Tatsumi Imada, Haruka Teraki, Kenta Miyoshi, Teppei Niide, Shori Inoue, Shogo Yoshimoto, Katsutoshi Hori, Hiroshi Shimizu, Yoshihiro Toya

## Abstract

*Acinetobacter* sp. strain Tol 5 is an attractive microorganism because of its ability to utilize various aromatic compounds, including toluene. However, because Tol 5 is a non-model microorganism, the utilization of carbon sources in central carbon metabolism remains unclear. In this study, the activity of the central carbon metabolism was characterized using ^13^C-metabolic flux analysis (^13^C-MFA). Tol 5 was cultured in synthetic medium using ethanol or acetate as the sole carbon source. In addition, because toluene is degraded into pyruvate and acetyl-CoA via the toluene dioxygenase pathway, this situation was mimicked by culturing with ethanol and lactate as co-substrates. The ^13^C-MFA confirmed flux through the glyoxylate shunt, which bypasses part of the TCA cycle when either ethanol or acetate was used as the sole carbon source. In ethanol, the flux ratio of isocitrate dehydrogenase at the isocitrate branch point decreased markedly. Unlike acetate, the conversion of ethanol to acetyl-CoA is accompanied by NADH production. Therefore, the observed change in this flux ratio was influenced by the excess NADH generated. Under ethanol-lactate conditions, Tol 5 simultaneously utilized ethanol and lactate. The ethanol consumption rate decreased to less than half of that observed when ethanol was the sole carbon source. Most of the lactate uptake was converted to pyruvate and metabolized via the TCA cycle. The glyoxylate shunt flux was lower than that in ethanol. These metabolic characteristics provide valuable insights into the use of Tol 5 as a next-generation microbial cell factory.

## Introduction

While microbial chemical production is attracting attention, there is a growing demand for utilizing not only sugars but also aromatic compounds that were previously difficult to use, such as benzene and toluene, which are petroleum-derived by-products. *Acinetobacter* sp. Tol 5 (Tol 5) is a gram-negative bacterium that was isolated from a toluene-removal biotrickling filter and can use various aromatic compounds, such as benzene, toluene, xylene, phenol, and stilbene (1). In addition, Tol 5 expresses a nanofiber protein called AtaA on its cell surface, allowing it to adhere to various carriers, such as glass and polyurethane, with high efficiency. Cells immobilized on carriers by AtaA are not easily detached, even during agitation of the culture medium. Immobilized cells on polyurethane foam can be repeatedly used as whole-cell biocatalysts (2). Cell immobilization using this method has also been applied to the reaction that converts indole into indigo (3). Thus, Tol 5 has attracted attention as a novel host microorganism for bioproduction because of its unique ability to utilize aromatic carbon sources and adhere to carrier surfaces.

Recently, the complete genome sequence of Tol 5 has been elucidated (4), and a genome editing technique has been established (5), leading to growing momentum for rational metabolic engineering. Investigations of metabolic pathways, such as the degradation pathways of aromatic compounds in these cells, have progressed (6). However, the regulation of metabolite flow in the central metabolism, which is crucial for precursor supply, remains unclear. Notably, the central carbon metabolism in *Acinetobacter* species has been suggested to differ from that of *E. coli*, as exemplified by the reported absence of hexokinase and pyruvate kinase (7,8). The ^13^C-metabolic flux analysis (^13^C-MFA) is a powerful method for revealing the intracellular metabolic flow in pathways (9,10). For this analysis, cells are cultured on substrates labeled with ^13^C at specific carbon atom positions. The uptake substrates are metabolized through various pathways, and the flux distribution is recorded as the ^13^C-labeling pattern of metabolites. The mass isotopomer distributions (MIDs) of the metabolites are measured using mass spectrometry, and the flux distribution is estimated by optimization to account for these measurements. To date, no study has investigated the metabolism of Tol 5 using ^13^C-MFA.

This research focused on detailing the carbon flow in the central carbon metabolism of Tol 5 grown in liquid culture. Because aromatic compounds enter the central carbon metabolism through long, complicated degradation steps, the use of a substrate that enters the central carbon metabolism more directly was considered to simplify the analysis. Tol 5 can be grown using aromatic compounds and ethanol (1,6). Ethanol is converted into acetate, which then enters the central carbon metabolism as acetyl-CoA (AcCoA). First, the flux distribution was characterized using ethanol or sodium acetate as the sole carbon source. In addition, it has been revealed that toluene is degraded through the toluene dioxygenase pathway and enters the central carbon metabolism as pyruvate and AcCoA (11). Toluene is poorly soluble in water and readily volatilizes, making it difficult to measure its specific consumption rate accurately. Among the available carbon sources of Tol 5, ethanol connects to AcCoA, and lactate connects to pyruvate. To investigate the pyruvate utilization pathways involved in toluene assimilation, the flux distribution was evaluated when ethanol and lactate were used as co-substrates.

## Materials and Methods

### Strain and medium

*Acinetobacter* sp. Tol 5 4140, an *ataA*-deletion mutant lacking the fiber protein AtaA, was used in this study (12). The contents of the Luria–Bertani (LB) medium and basal salt (BS) medium have been previously described (1). Ethanol (30 mM), sodium acetate (30 mM), or both ethanol and lactate (15 mM each) were used as carbon sources.

### Culture conditions

Cells were inoculated from glycerol stock into 5 mL of LB medium in a test tube and incubated at 28°C with 115 rpm overnight. For preculture, cells from the starter were collected by centrifugation and washed twice with BS medium lacking trace metals. Subsequently, cells were inoculated into 20 mL of BS medium to an optical density at 600 nm (OD_600_) of 0.1 in a 100-mL flask, followed by overnight shaking at 28°C with 115 rpm. The main culture was inoculated in the same manner as the preculture and incubated at 28°C with 115 rpm. Cultures were performed in triplicate.

### Measurement of dry cell weight and extracellular metabolites

The cell concentration was measured as OD_600_ using a UVmini-1240 (Shimadzu Corporation, Japan). The conversion factor between OD_600_ and dry cell weight (DCW) was determined as follows: A culture volume that satisfied OD_600_ × culture volume (mL) = 25 was collected and filtered under vacuum using a membrane filter (H050A090C, Advantec Toyo). After thoroughly drying for more than two days in a dryer at 60°C, the weight of the filter was measured, and the previously measured weight of the filter was subtracted to calculate the DCW. The concentrations of acetate,ethanol, and lactate in the culture supernatant were measured using high performance liquid chromatography (HPLC) Prominence (Shimadzu). The detailed analytical conditions have been described in a previous study (13).

### Measurement of ^13^C-enrichment of proteinogenic amino acids

Cells were collected during the exponential growth phase at around OD_600_ ≅ 1, by centrifugation (3,700×g, 4°C, 15 min) to obtain an amount satisfying OD_600_ × culture volume (mL) = 3. After washing the cells with 0.9% NaCl, the cells were suspended in 1 mL of 6 N HCl and hydrolyzed at 105°C for 18 h. Proteinogenic amino acids were derivatized with *tert*-butyldimethylsilyl (TBDMS) and analyzed by gas chromatography mass spectrometry (GC/MS). The detailed pretreatment and analytical conditions have been described in a previous study (13).

### ^13^C-MFA

Central carbon metabolism was reconstructed, including the TCA cycle, glyoxylate shunt, gluconeogenesis pathway, pentose phosphate pathway, based on the genomic information of Tol 5 (Fig. 1). The reactions and carbon atom transitions of the central carbon metabolism and amino acid synthesis are listed in Table S1 and S2, respectively. A model for calculating the MIDs of amino acids from the flux distributions and substrate label patterns was constructed based on an elementary metabolite unit framework (14). The precursor requirements for biomass formation were taken from the analysis of closely related strain *Pseudomonas putida* (15). Metabolic fluxes were estimated by minimizing the residual sum of squares (RSS) between the measured and estimated MIDs of the proteinogenic amino acids and effluxes (Eq. 1), and the goodness of fit was evaluated using a chi-square test.

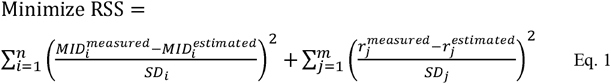

where 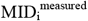 and 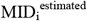 are the measured and estimated MID of *i*th amino acid, 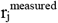 and 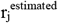 are the measured and estimated efflux of *j*th reaction, respectively. SD_i_ and SD_j_ are the standard deviations (SD) of MID and efflux, respectively. The SD of MIDs was set to 0.01 (13), and the SD of effluxes was calculated from replicate measurements. The 95% confidence intervals for the estimated fluxes were calculated using a grid search algorithm (16). All calculations for MID correction (natural isotope abundance and carry-over from the inoculum) and ^13^C-MFA were performed using OpenMebius2 (17).

**Fig. 1.**
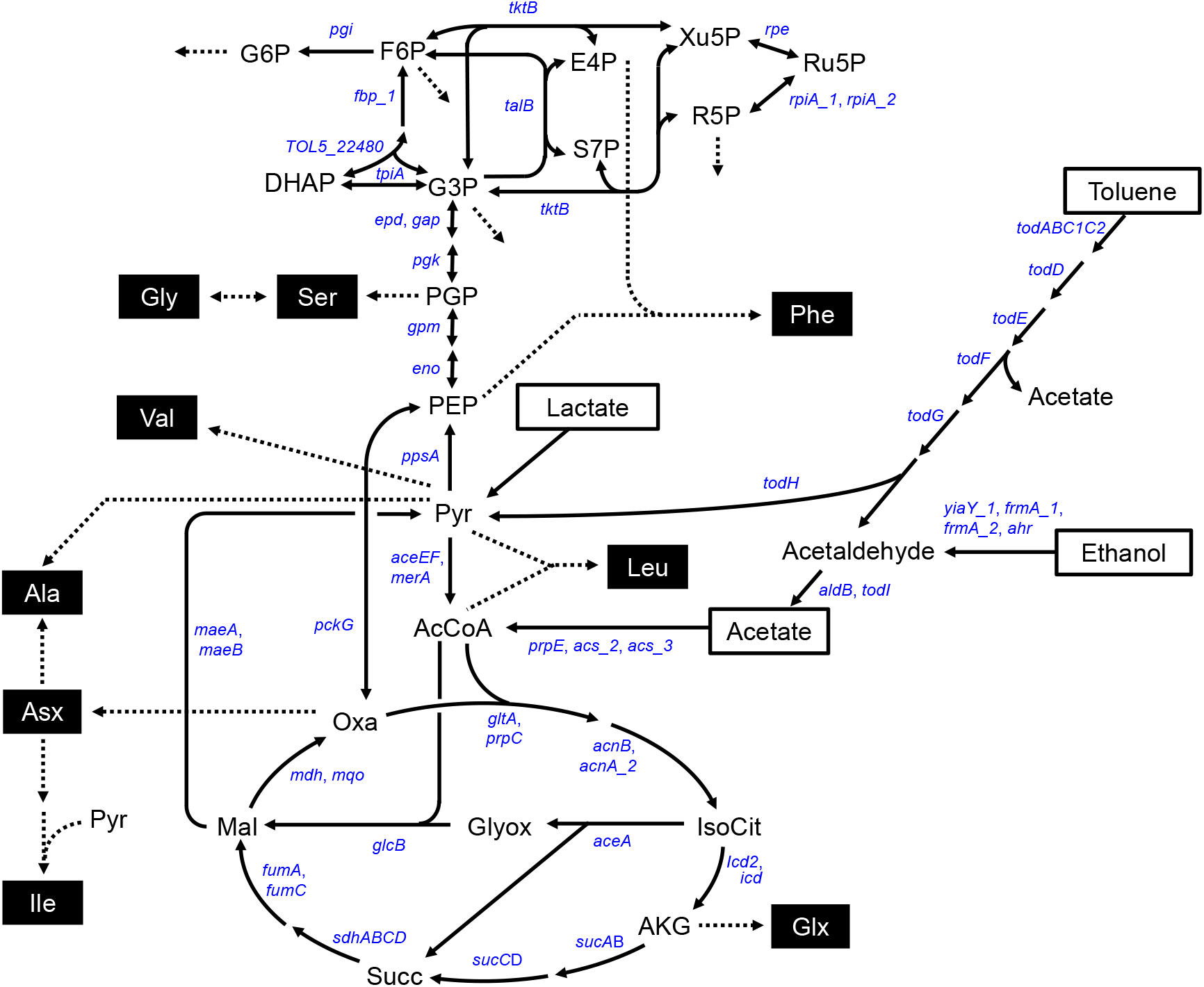
Central carbon metabolism of Tol 5 for ^13^C-MFA. Flux estimation was performed using the MIDs of amino acids depicted in boxes with white text on a black background. Carbon sources are shown in boxes with black text on a white background. Dotted lines indicate biosynthetic pathways including amino acid syntheses. Gene names are shown in italics. Abbreviations of metabolites: AcCoA, acetyl-CoA; AKG, 2-ketoglutarate; DHAP, dihydroxyacetone phosphate; E4P, erythrose 4-phosphate; F6P, fructose 6-phosphate; G3P, glyceraldehyde 3-phosphate; G6P, glucose 6-phosphate; Glyox, glyoxylate; IsoCit, isocitrate; Mal, malate; PEP, phosphoenolpyruvate; PGP, 3-phosphoglycerate; Pyr, pyruvate; S7P, sedoheptulose 7-phosphate; Succ, succinate; R5P, ribose 5-phosphate; Ru5P, ribulose 5-phosphate; Xu5P, xylulose 5-phosphate.

## Results

### Assessment of carbon source utilization of Tol 5

First, Tol 5 was cultured with ethanol or sodium acetate as the sole carbon source to evaluate the effect of carbon flow on metabolism. In Tol 5, ethanol is converted to acetate via acetaldehyde and incorporated into the central metabolism as AcCoA. The wild-type strain of Tol 5 highly aggregated in liquid culture owing to the presence of the nanofiber protein AtaA, making it difficult to measure the OD_600_. Therefore, a strain lacking *ataA* was used to facilitate handling in a liquid culture. The growth profiles are shown in Fig. 2. HPLC analysis of the culture supernatants did not detect any metabolites other than the substrates. The specific growth rate in ethanol was 25% higher than that in acetate (Table 1). DCW was required to calculate the specific substrate consumption rates from the time course of OD_600_ and substrate concentration.

**Table 1.** Specific rates of Tol 5 under different carbon sources.

| | Growth rate | Consumption rate ($\text{mmol g}^{-1} \text{h}^{-1}$) | | |
| --- | --- | --- | --- | --- |
| Carbon source | ($\text{h}^{-1}$) | Acetate | Ethanol | Lactate |
| Acetate | $0.42 \pm 0.06$ | $17.9 \pm 0.9$ | - | - |
| Ethanol | $0.53 \pm 0.01$ | - | $16.3 \pm 3.8$ | - |
| Ethanol + Lactate | $0.60 \pm 0.02$ | - | $6.9 \pm 2.5$ | $9.7 \pm 1.3$ |

**Fig. 2.**
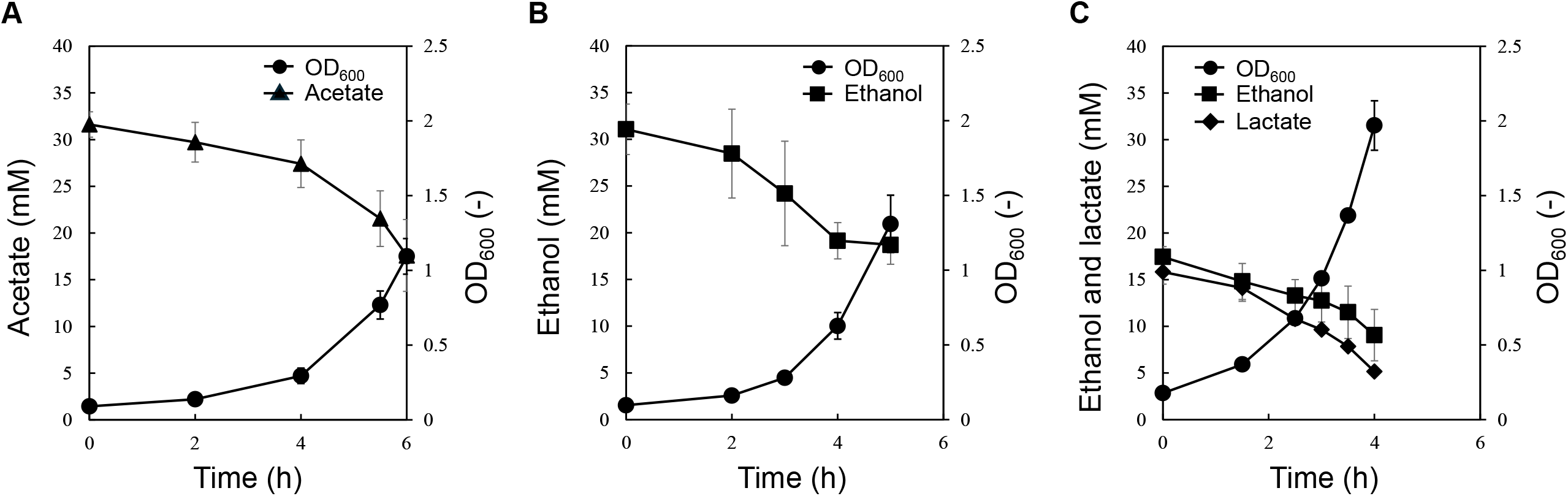
Growth profiles of Tol 5 under different carbon sources. (A) acetate, (B) ethanol, and (C) ethanol and lactate as carbon sources. Filled cycles, squares, triangles, and diamonds represent OD_600_ and the concentration of acetate, ethanol, and lactate, respectively.

The measured conversion factor between OD_600_ and DCW was 0.40 ± 0.02 g L^-1^ OD_600_ ^-1^ (n = 5). Table 1 presents the specific consumption rates. No significant differences were observed in the specific consumption rates of acetate or ethanol. The higher growth rate observed in the presence of ethanol may have been due to NADH production during the conversion of ethanol to acetate.

### Central carbon metabolism and amino acid synthesis pathways of Tol 5

Knowledge of metabolic pathways is required to investigate the mechanism by which the assimilated carbon source is metabolized and its flux distribution. For Tol 5, genomic information was elucidated, and its metabolic pathways were illustrated in the KEGG database. Fig. S1 compares the enzymes present in *Escherichia coli* and Tol 5 in the main pathways. Tol 5 lacks pyruvate kinase and phosphofructokinase in glycolysis but possesses other enzymes similar to *E. coli*. Consistent with the absence of a complete glycolysis pathway, Tol 5 has been reported not to grow on sugars as sole carbon sources, unlike *E. coli* (6).

To perform ^13^C-MFA in growing cells, labeling patterns of proteinogenic amino acids are generally employed. The amino acid biosynthesis pathways in Tol 5 were also compared with those in *E. coli*. As shown in Fig. S1, although most biosynthesis pathways are similar, *E. coli* synthesized alanine only from pyruvate, whereas Tol 5 synthesized alanine from both pyruvate and aspartate.

### Analysis of ^13^C-enrichment of proteinogenic amino acids

The ^13^C-labeling experiments were conducted to reveal carbon flow in the pathways in detail. Flux distribution information is reflected in the ^13^C-labeling patterns of proteinogenic amino acids. In this study, [1-^13^C] ethanol or sodium acetate was used as the carbon source. These substrates consist of two carbon atoms; therefore, labeling either carbon atom is equivalent. Proteinogenic amino acids were obtained from cells in the exponential growth phase and measured by GC/MS following TBDMS derivatization. The MIDs of the fragment ions ([M-57]^+^, [M-85]^+^, [M-159]^+^, and f302^+^) of the derivatized amino acids were detected. MIDs corrected for natural isotope abundance and carry-over from the inoculum are shown in Table S3. The amino acid synthesis pathways are shown in Fig. 1.

A scatter plot of the MIDs in ethanol and acetate is shown in Fig. 3A. The differences in the MIDs suggest differences in the flux distributions. Among the amino acids, the MIDs of Asx and Thr synthesized from oxaloacetate (Oxa) differed markedly between ethanol and acetate (Fig. 3B). Although the M+1 and M+2 ratios of the MIDs were almost identical in acetate, the M+2 ratio was dominant in ethanol. Acetate and ethanol are converted to AcCoA and then incorporated into the TCA cycle. Isocitrate is subsequently catalyzed by isocitrate dehydrogenase (ICDH) and isocitrate lyase (ICL). Because the metabolites through these two branched routes merge into Oxa, the MIDs of the amino acids derived from Oxa contain the flux ratio information between the TCA cycle and the glyoxylate shunt.

**Fig. 3.**
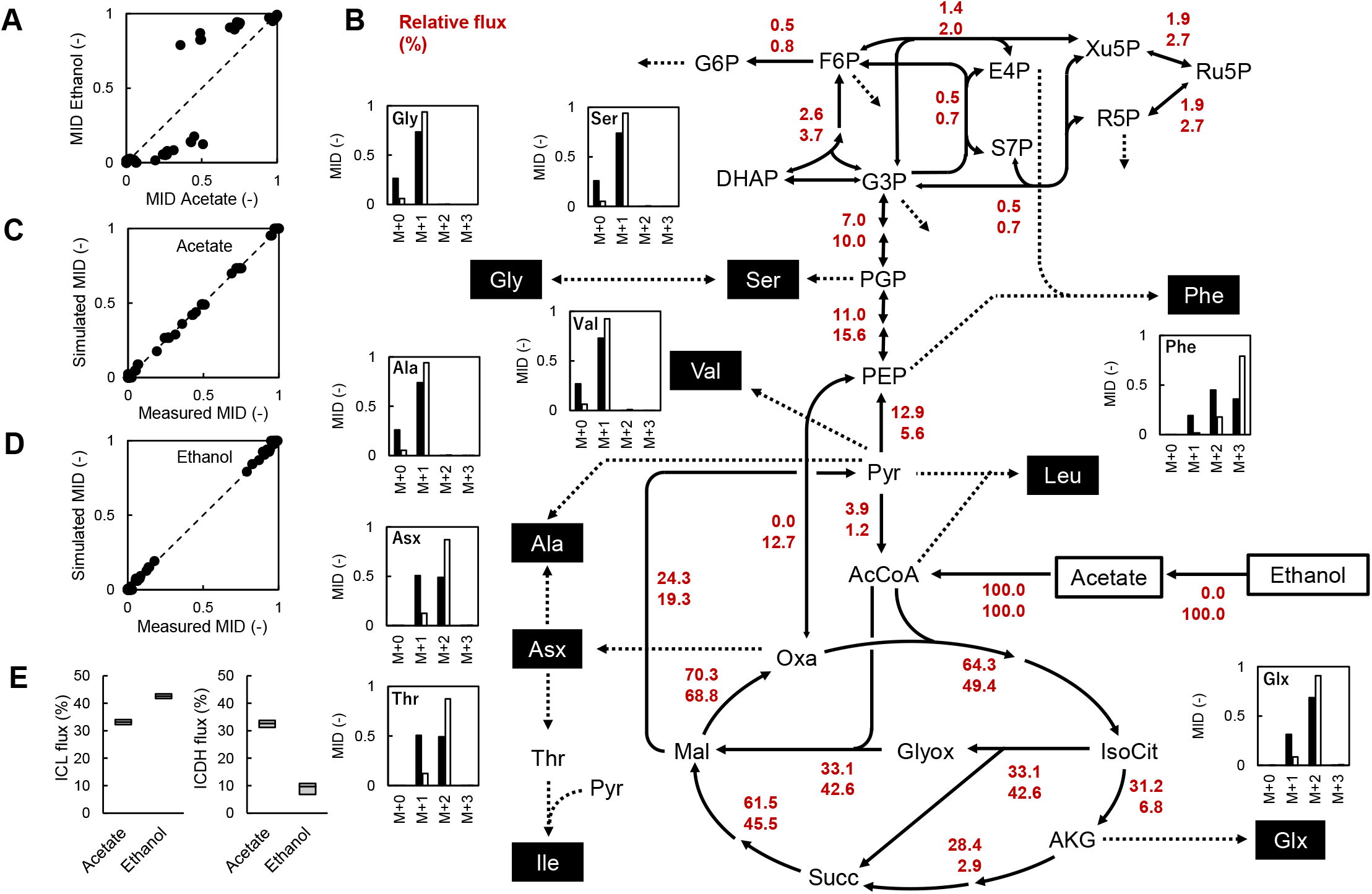
Metabolic flux distributions of Tol 5 under acetate and ethanol conditions. (A) Scatter plot of amino acid MIDs under acetate (x-axis) and ethanol (y-axis) conditions. (B) Flux distributions. Upper and lower values represent fluxes under the acetate and ethanol conditions, respectively. Fluxes were normalized to an acetate or ethanol uptake rate of 100. Dashed arrows represent efflux to cellular biosynthesis, including amino acid syntheses. Bar graphs show the relative abundance of amino acid MID (measured) from M+0 to M+3 of the [M-57]^+^, which contains all skeletal carbons. Filled and open bars indicate the acetate and ethanol conditions, respectively. Ile and Leu are not shown because their [M-57]^+^ fragments could not be measured. (C, D) Scatter plot of measured versus simulated MIDs under the acetate (C) and ethanol (D) conditions. (E) 95% confidence interval of the estimated fluxes. Box indicates the confidence interval range, and vertical lines indicates the optimal value. Abbreviations are as in Fig. 1.

### Flux distributions under acetate or ethanol conditions

The flux distribution was estimated using a model to explain the measured MIDs. Optimization was performed to minimize the RSS between the measured and simulated MIDs from the estimated flux distribution. The threshold of the chi-square test for the goodness of fit was 104.1. The RSSs under acetate and ethanol conditions were 94.4 and 148.9, respectively. The flux distribution under the acetate condition passed the chi-square test, whereas that under the ethanol condition did not. However, the simulated MIDs agreed equally well with the measured MIDs in both conditions (R^2^ = 0.999; Fig. 3C and 3D). A comparison between the measured and simulated MIDs is summarized in Table S3. The estimated flux distributions under the acetate and ethanol conditions are shown in Fig. 3B. Each flux value was normalized so that the acetate or ethanol uptake rate was 100. The 95% confidence intervals of all estimated fluxes are listed in Table S4 (acetate condition) and Table S5 (ethanol condition).

A comparison between acetate and ethanol conditions suggested that the proportion of isocitrate metabolized via the TCA cycle through 2-ketoglutarate (AKG) was significantly lower under ethanol conditions. The ICL:ICDH flux ratios at the isocitrate branch point were 51:49 under acetate conditions, whereas they were 86:14 under ethanol conditions. To confirm the differences in the fluxes, 95% confidence intervals of the flux, which statistically accounted for the measured MIDs, were compared (Fig. 3E). The confidence intervals of ICDH did not overlap between conditions, indicating that this flux was significantly lower in ethanol.

The flux from AcCoA and Oxa to isocitrate was higher under acetate conditions because AcCoA was also used for the glyoxylate shunt. Although differences were observed in the flux of the route from malate to phosphoenolpyruvate (PEP), the fluxes of the route through pyruvate or Oxa could not be distinguished, and there was no difference in carbon atom transitions. The overall flux distribution of gluconeogenesis was similar in the acetate and ethanol conditions.

### Flux distribution under ethanol and lactate co-utilization

When toluene is degraded in Tol 5, it is incorporated into central carbon metabolism as both pyruvate and AcCoA. Among the available carbon sources, ethanol is connected to AcCoA and lactate to pyruvate. To mimic toluene assimilation, Tol 5 was cultured with ethanol and lactate as co-substrates (Fig. 2C). The culture profile suggested that Tol 5 consumed ethanol and lactate simultaneously. The specific rates during the exponential phase are summarized in Table 1. Under ethanol-lactate conditions, the specific growth rate increased by approximately 10% relative to ethanol-only conditions, whereas the specific ethanol consumption rate decreased by almost half. In addition, the consumption rate of lactate was higher than that of ethanol under the co-substrate conditions.

To confirm the contribution of carbon atoms derived from ethanol, cells were cultured with [U-^13^C] ethanol and unlabeled lactate. The MIDs of the proteinogenic amino acids are shown in Fig. S2. Because unlabeled lactate was incorporated into pyruvate, Ala and Val synthesized from pyruvate did not contain ^13^C atoms. In contrast, the ^13^C atoms were incorporated into Ser, Gly, and Phe which were synthesized from the upstream glycolytic intermediates PEP and 3PG. This indicates that carbon atoms derived from ethanol are converted to PEP via the TCA cycle and PEP carboxykinase and were used in gluconeogenesis.

For ^13^C-MFA, cells were cultured using [1-^13^C] ethanol and unlabeled lactate. Flux distribution was estimated using the measured MIDs (Table S3). The estimated flux distribution under ethanol-lactate conditions is shown in Fig. 4A. To allow a comparison with the fluxes under ethanol-only conditions, the reaction fluxes are shown as absolute values (mmol g^-1^ h^-1^). Because the RSS was 82.1 (the threshold of the chi-square test was 103.0), a flux distribution that consistently accounted for the experimental data was estimated (Fig. 4B). The 95% confidence intervals of all estimated fluxes are listed in Table S6.

**Fig. 4.**
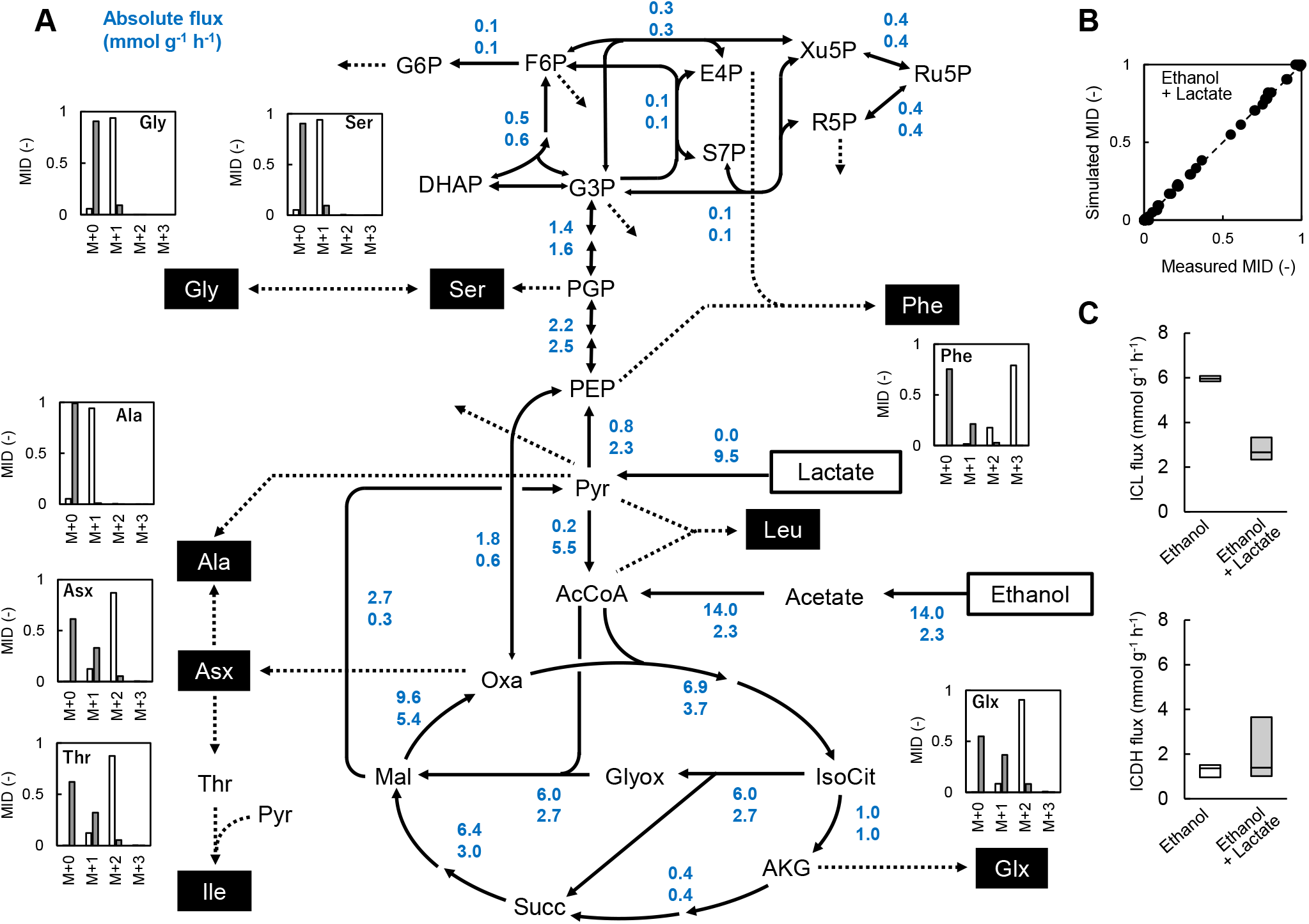
Metabolic flux distributions of Tol 5 under ethanol or ethanol–lactate conditions. (A) flux distributions. The upper and lower values represent absolute flux values (mmol g^-1^ h^-1^) under ethanol-only and ethanol-lactate conditions, respectively. The bar graphs show the relative abundance of amino acid MID from M+0 to M+3 for the [M-57]^+^ fragment, which contains all skeletal carbons. The right and left bars represent the ethanol-only and ethanol-lactate conditions, respectively. (B) Scatter plot of measured versus simulated MIDs under the ethanol–lactate conditions. (C) 95% confidence interval of the estimated flux. Box indicates the confidence interval range, and the vertical line indicates the optimal value. Abbreviations are as in Fig. 1.

After entering the central carbon metabolism as pyruvate, lactate is metabolized through two branches, one leading to PEP and the other to AcCoA. Pyruvate flux was distributed 24% to PEP and 58% to AcCoA. The existence of lactate-derived AcCoA was consistent with the observation that amino acids synthesized via the TCA cycle, such as Glx, Asx, and Thr, contain fewer ^13^C atoms than those synthesized under ethanol-only conditions. The flux through the glyoxylate shunt was lower under ethanol-lactate conditions than under ethanol-only conditions. The ICL:ICDH flux ratio at the isocitrate branch point was 73:27 under ethanol-lactate conditions and 86:14 under ethanol conditions (Fig. 4A and 4C).

## DISCUSSION

In the present study, the intracellular metabolic phenotypes of Tol 5 with different carbon sources were investigated using ^13^C-labeling techniques. The flux distribution during the exponential growth phase was determined under acetate-only, ethanol-only, or ethanol-lactate conditions. Both acetate and ethanol enter central carbon metabolism as AcCoA, and no significant difference was observed in their consumption rates under acetate- and ethanol-only conditions. Since NADH is produced when ethanol is converted to acetate, NADH production appears to be higher under ethanol conditions than under acetate conditions. In addition, when growing using only C2 compounds, such as acetate, as the sole carbon source, the use of a glyoxylate shunt is essential (18). The glyoxylate shunt bypasses the CO_2_-emitting reactions in the TCA cycle, allowing carbon to be used for biomass building block formations. A certain amount of glyoxylate shunt flux was observed under both acetate- and ethanol-only conditions.

The regulatory mechanism controlling the flux ratio between the TCA cycle and the glyoxylate shunt has been well studied in *E. coli* (19). ICDH has a lower affinity constant (Km) for isocitrate than ICL; therefore, isocitrate naturally tends to flow toward the TCA cycle (20). *E. coli* regulates the flux ratio at the isocitrate branch point by phosphorylating ICDH (21). Under acetate conditions, the expression of the *aceBAK* operon increased, and the ICDH kinase activity of the regulatory enzyme AceK was enhanced. Consequently, ICDH activity decreases owing to its phosphorylation, redirecting the metabolic pathway to the glyoxylate shunt (22). The glyoxylate shunt flux is almost negligible under glucose batch conditions in *E. coli* (23), whereas approximately 30% of the isocitrate flows through the glyoxylate shunt under acetate continuous culture conditions (24). In Tol 5, 51% of the isocitrate flowed through the glyoxylate shunt under acetate conditions, and the contribution of the glyoxylate shunt was higher than that of *E. coli*. Although no protein with an amino acid sequence similar to *E. coli* AceK was found in the genome of Tol 5, it has been reported that ICDH activity is regulated by phosphorylation in the related species *Acinetobacter baumannii* (25), suggesting the presence of a regulatory protein with an AceK-like function in *Acinetobacter* species. Another report indicated that the expression of *aceA*, which encodes ICL, depends on acetate concentration in *Acinetobacter oleivorans* DR1 (26). To determine the regulatory mechanism of ICDH inhibition or ICL activation under acetate conditions in Tol 5, it is necessary to evaluate the enzyme activities of ICDH and ICL, the phosphorylation level of ICDH, and the Km values of ICDH and ICL for isocitrate.

The flux ratios at the isocitrate branch point differed between the acetate and ethanol conditions in Tol 5, with a marked reduction in ICDH flux observed under ethanol conditions (Fig. 3B and 3C). The ICDH reaction generates the reducing power of NADPH (25). Because two molecules of NADH are generated during the conversion of one molecule of ethanol to one molecule of acetate, the low ICDH flux observed under ethanol conditions would prevent the excess production of reducing power. In the *pgi* deletion mutant of *E. coli*, the use of the oxidative pentose phosphate pathway is enforced, resulting in excess NADPH generation. To prevent further NADPH generation by ICDH, TCA cycle flux is redirected to the glyoxylate shunt (27). Similarly, to alleviate NADPH generation under ethanol conditions, Tol 5 may regulate the flux ratio to bypass ICDH reaction. One question is why the redox cofactor generated in ICDH is NADPH, whereas that generated during a conversion of ethanol to acetate is NADH. In *Acinetobacter* sp. 1NG, the specific enzymatic activity of ICDH was high whereas that of AKG dehydrogenase was low under ethanol conditions (28). Because AKG dehydrogenase is involved in the generation of NADH, the inhibition of this reaction by NADH may have switched the metabolic flow from the TCA cycle to the glyoxylate shunt.

The advantage of Tol 5 is its ability to use aromatic compounds such as toluene, which are typically challenging for common hosts such as *E. coli* to use (1). Recent genetic research indicates that Tol 5 breaks down toluene into pyruvate and AcCoA via the toluene dioxygenase pathway (11). However, its function in the central carbon metabolism remains unclear. The ^13^C-MFA is a powerful approach for uncovering pathway flux distribution, although it requires accurate input and output flux measurements. Because of the challenges in accurately measuring the specific consumption rate of toluene attributed to its volatility and limited water solubility, we mimicked toluene assimilation using lactate and ethanol as co-substrates. Tol 5 simultaneously consumed lactate and ethanol, allowing the determination of the flux distribution. The ethanol consumption rate markedly decreased under ethanol-lactate conditions compared to ethanol-only conditions (Fig. 4A). AcCoA produced from lactate may inhibit the formation of AcCoA from ethanol. Under conditions with both ethanol and lactate, the glyoxylate shunt flux was reduced compared to that when only ethanol was present. Under ethanol-only conditions, the use of the glyoxylate shunt was essential to supply biomass building block precursors to bypass the CO_2_ emission reactions in the TCA cycle; however, under ethanol-lactate conditions, these precursors can be supplied from lactate, so the necessity of the glyoxylate shunt decreased. In addition, a reduction in the specific ethanol consumption rate led to a decrease in NADH production, thereby alleviating the inhibition of the TCA cycle downstream of ICDH. Thus, the flux distribution obtained under the ethanol-lactate condition may provide a useful reference for future comparison with actual aromatic-compound assimilation. These observations also suggest that co-substrate feeding could be explored as a strategy to modulate central carbon allocation in Tol 5.

The estimated flux distributions provide insight into metabolic engineering to produce valuable compounds using Tol 5, a potential host for next-generation microbial cell factory. For example, to increase the production of a target compound, we can determine the reactions that should be enhanced or suppressed through simulations using the actual flux distribution (29). The appeal of Tol 5 lies in its adhesive nanofiber proteins that allow it to be strongly immobilized on carriers. A recent study demonstrated that Tol 5 exhibits desiccation tolerance, and that immobilized cells can degrade toluene in the gas phase (30). However, its metabolic phenotype under immobilized or gas-phase conditions remains largely unknown. Future comparison of the present liquid-culture flux maps with those obtained under immobilized or gas-phase conditions may reveal how these distinct environments reshape central carbon metabolism during bioprocess operation. Thus, the flux distributions established here provide quantitative reference states for understanding how the metabolism of Tol 5 changes in response to different substrates and conditions.

## CRediT authorship contribution statement

**Erina Miyamoto:** Data curation, Formal analysis, Investigation. **Tatsumi Imada:** Formal analysis, Investigation, Software, Validation, Writing – review & editing. **Haruka Teraki:** Formal analysis. **Kenta Miyoshi:** Formal analysis. **Teppei Niide:** Formal analysis. **Shori Inoue:** Formal analysis. Writing – review & editing. **Shogo Yoshimoto:** Conceptualization, Writing – review & editing. **Katsutoshi Hori:** Conceptualization, Resources, Project administration, Writing – review & editing. **Hiroshi Shimizu:** Conceptualization, Supervision, Writing – review & editing. **Yoshihiro Toya:** Conceptualization, Formal analysis, Investigation, Methodology, Visualization, Writing – original draft.

## Acknowledgements

We thank Dr. Hiroshi Tsugawa (Kyoto University), Mr. Taisei Naobayashi and Ms. Kanako Tokiyoshi (Tokyo University of Agriculture and Technology) for useful discussion. This work was supported by a Grant-in-Aid for Scientific Research (S) No. JP25K24646 from the Japan Society for the Promotion of Science (JSPS), the Green Technologies of Excellence (GteX) Program Japan Grant Number JPMJGX23B4, the Adopting Sustainable Partnerships for Innovative Research Ecosystem (ASPIRE) Grant Number JPMJAP24A2, and K Program Grant Number JPMJKP25V2 from the Japan Science and Technology Agency (JST), and JP23wm0325066 from the Japan Agency for Medical Research and Development (AMED).

