## Supplemental data for "Characterization of central carbon metabolism of *Acinetobacter* sp. Tol 5 based on ^13^C-metabolic flux analysis"

**Miyamoto *et al.***

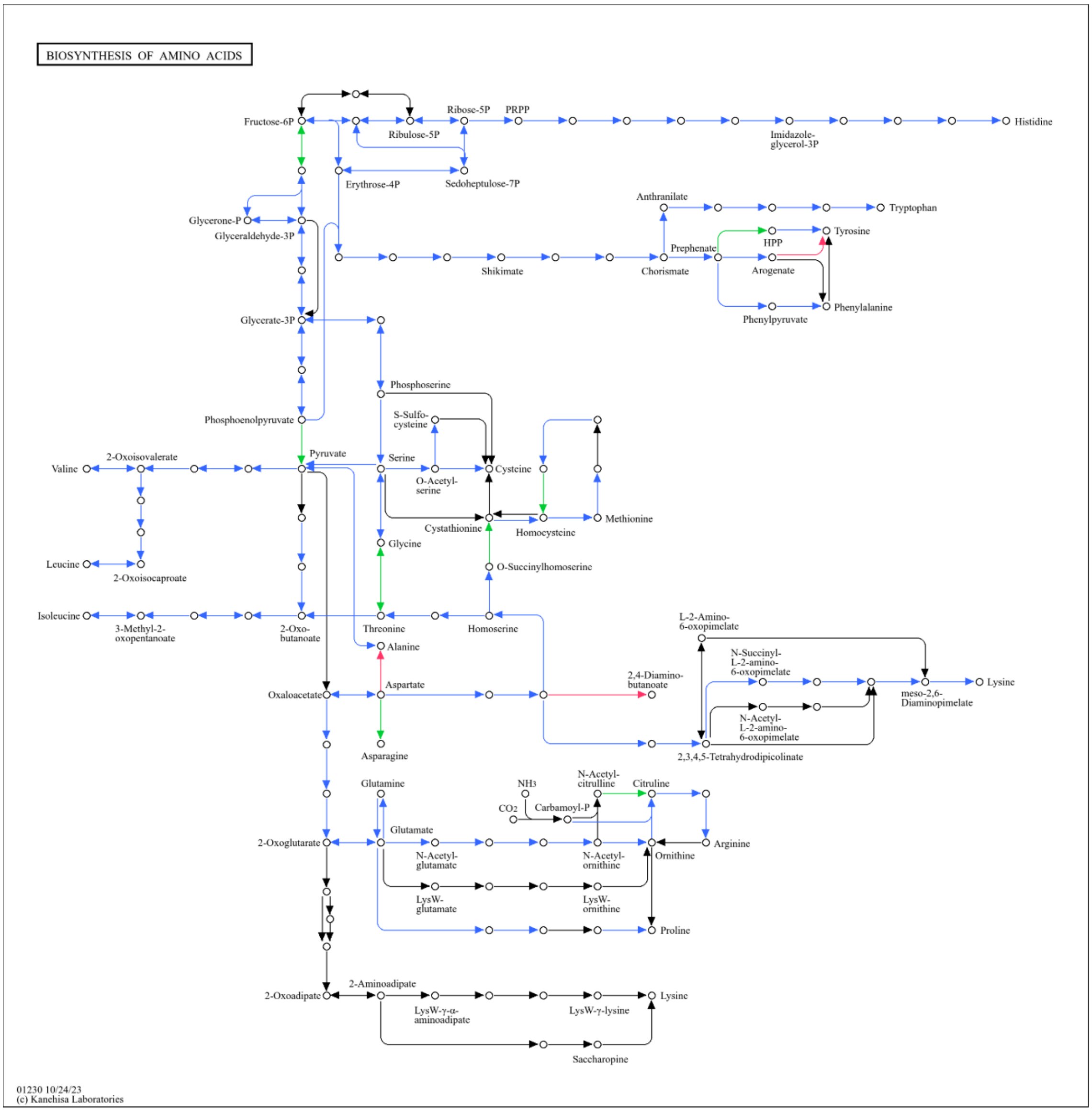


**Fig. S1.** Differences in amino acid biosynthesis pathways between E. coli and Tol 5 on the KEGG pathway map.

Blue arrows indicate reactions shared by both *E. coli* and Tol 5. Green and red arrows indicate reactions present only in *E. coli* and only in Tol 5, respectively.

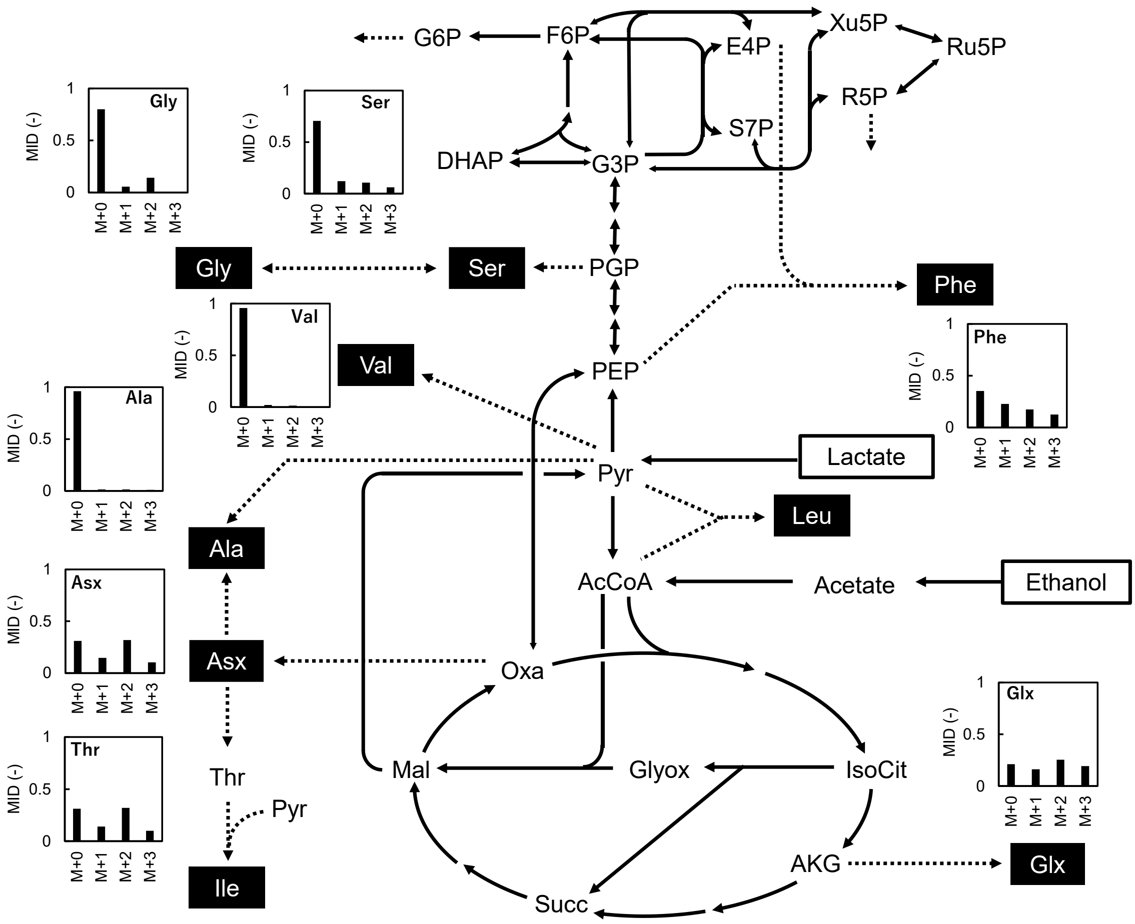


**Fig. S2.** MIDs of proteinogenic amino acids in Tol 5 cultured with [U-$^{13}$C] ethanol and unlabeled lactate

The bar graphs show the relative abundance of amino acid MID from M+0 to M+3 for the [M-57]$^{+}$ fragment, which contains all skeletal carbons.

**Table S1.** Metabolic model.

| ID | Reaction | Carbon transition |
|---|---|---|
| r1 | Subs_C2 --> AcCoA | AB --> AB |
| r2 | AcCoA + OAA --> IsoCit | AB + CDEF --> FEDBAC |
| r3 | IsoCit --> AKG + CO2_in | ABCDEF --> ABCDE + F |
| r4 | AKG --> Sym_SUC + CO2_in | ABCDE --> BCDE + A |
| r5 | Sym_SUC <=> MAL | ABCD --> ABCD |
| r6 | MAL <=> OAA | ABCD --> ABCD |
| r7 | IsoCit + AcCoA --> MAL + Sym_SUC | ABCDEF + GH --> ABHG + FCDE |
| r8 | PEP + CO2_in <=> OAA | ABC + D --> ABCD |
| r9 | MAL --> PYR + CO2_in | ABCD --> ABC + D |
| r10 | Subs_C3 --> PYR | ABC --> ABC |
| r11 | PYR --> PEP | ABC --> ABC |
| r12 | PEP <=> PGP | ABC --> ABC |
| r13 | PGP <=> G3P | ABC --> ABC |
| r14 | G3P <=> DHAP | ABC --> ABC |
| r15 | DHAP + G3P <=> FBP | CBA + DEF --> ABCDEF |
| r16 | FBP --> F6P | ABCDEF --> ABCDEF |
| r17 | F6P <=> G6P | ABCDEF --> ABCDEF |
| r18 | PYR --> AcCoA + CO2_in | ABC --> BC + A |
| r19 | Ru5P <=> X5P | ABCDE --> ABCDE |
| r20 | Ru5P <=> R5P | ABCDE --> ABCDE |
| r21 | R5P + X5P <=> S7P + G3P | ABCDE + FGHIJ --> FGABCDE + HIJ |
| r22 | G3P + S7P <=> F6P + E4P | ABC + DEFGHIJ --> DEFABC + GHIJ |
| r23 | E4P + X5P <=> F6P + G3P | ABCD + EFGHI --> EFABCD + GHI |
| r24 | PGP --> Ser | ABC --> ABC |
| r25 | Ser <=> Gly + THF_in | ABC <=> AB + C |
| r26 | OAA --> ALA + CO2_in | ABCD --> DCB + A |
| r27 | PYR --> ALA | ABC --> ABC |
| r28 | Subs_CO2 <=> CO2_in | A --> A |
| r29 | Subs_THF <=> THF_in | A --> A |

**Table S2.** Carbon transitions in amino acid synthesis.

| Metabolite | Reaction | Carbon transitions |
|---|---|---|
| Ala57 | ALA --> Ala57 | ABC --> ABC |
| Ala85 | ALA --> Ala85 | ABC --> BC |
| Asx57 | OAA --> Asx57 | ABCD --> ABCD |
| Asx85 | OAA --> Asx85 | ABCD --> BCD |
| Asx159 | OAA --> Asx159 | ABCD --> BCD |
| Asx302 | OAA --> Asx302 | ABCD --> AB |
| Glx57 | AKG --> Glx57 | ABCDE --> ABCDE |
| Glx85 | AKG --> Glx85 | ABCDE --> BCDE |
| Glx159 | AKG --> Glx159 | ABCDE --> BCDE |
| Gly57 | PGP --> Gly57 | ABC --> AB |
| Gly85 | PGP --> Gly85 | ABC --> B |
| Ile85 | OAA + PYR --> Ile85 | ABCD + EFG --> BFGCD |
| Ile159 | OAA + PYR --> Ile159 | ABCD + EFG --> BFGCD |
| Leu85 | AcCoA + PYR + PYR --> Leu85 | AB + CDE + FGH --> BDEGH |
| Leu159 | AcCoA + PYR + PYR --> Leu159 | AB + CDE + FGH --> BDEGH |
| Phe57 | E4P + PEP + PEP --> Phe57 | ABCD + EFG + HIJ --> EFGIJABCD |
| Phe85 | E4P + PEP + PEP --> Phe85 | ABCD + EFG + HIJ --> FGIJABCD |
| Phe159 | E4P + PEP + PEP --> Phe159 | ABCD + EFG + HIJ --> FGIJABCD |
| Phe302 | E4P + PEP + PEP --> Phe302 | ABCD + EFG + HIJ --> EF |
| Ser57 | Ser --> Ser57 | ABC --> ABC |
| Ser85 | Ser --> Ser85 | ABC --> BC |
| Ser159 | Ser --> Ser159 | ABC --> BC |
| Ser302 | Ser --> Ser302 | ABC --> AB |
| Val57 | PYR + PYR --> Val57 | ABC + DEF --> ABECF |
| Val85 | PYR + PYR --> Val85 | ABC + DEF --> BECF |
| Val159 | PYR + PYR --> Val159 | ABC + DEF --> BECF |

**Table S3.** Measured and estimated MIDs of amino acids.

| Name | Fragment | MID | Acetate | | Ethanol | | Ethanol + Lactate | |
|---|---|---|---|---|---|---|---|---|
| | | | Measured | Estimated | Measured | Estimated | Measured | Estimated |
| Ala | [M-57]+ | M + 0 | 0.259 | 0.267 | 0.053 | 0.060 | 0.989 | 0.989 |
| | | M + 1 | 0.740 | 0.733 | 0.941 | 0.940 | 0.010 | 0.011 |
| | | M + 2 | 0.000 | 0.000 | 0.005 | 0.000 | 0.000 | 0.000 |
| | | M + 3 | 0.001 | 0.000 | 0.001 | 0.000 | 0.000 | 0.000 |
| | [M-85]+ | M + 0 | 0.996 | 1.000 | 0.987 | 1.000 | 0.994 | 1.000 |
| | | M + 1 | 0.001 | 0.000 | 0.008 | 0.000 | 0.006 | 0.000 |
| | | M + 2 | 0.003 | 0.000 | 0.005 | 0.000 | 0.000 | 0.000 |
| Asx | [M-57]+ | M + 0 | 0.001 | 0.023 | 0.000 | 0.004 | 0.615 | 0.615 |
| | | M + 1 | 0.507 | 0.488 | 0.125 | 0.126 | 0.331 | 0.335 |
| | | M + 2 | 0.489 | 0.489 | 0.871 | 0.870 | 0.053 | 0.051 |
| | | M + 3 | 0.001 | 0.000 | 0.004 | 0.000 | 0.001 | 0.000 |
| | | M + 4 | 0.001 | 0.000 | 0.000 | 0.000 | 0.001 | 0.000 |
| | [M-85]+ | M + 0 | 0.246 | 0.267 | 0.054 | 0.074 | 0.779 | 0.780 |
| | | M + 1 | 0.752 | 0.733 | 0.938 | 0.926 | 0.219 | 0.220 |
| | | M + 2 | 0.001 | 0.000 | 0.006 | 0.000 | 0.001 | 0.000 |
| | | M + 3 | 0.001 | 0.000 | 0.001 | 0.000 | 0.001 | 0.000 |
| | [M-159]+ | M + 0 | 0.242 | 0.267 | 0.057 | 0.074 | 0.769 | 0.780 |
| | | M + 1 | 0.746 | 0.733 | 0.927 | 0.926 | 0.216 | 0.220 |
| | | M + 2 | 0.004 | 0.000 | 0.008 | 0.000 | 0.005 | 0.000 |
| | | M + 3 | 0.009 | 0.000 | 0.008 | 0.000 | 0.010 | 0.000 |
| | f302+ | M + 0 | 0.265 | 0.268 | 0.059 | 0.060 | 0.781 | 0.784 |
| | | M + 1 | 0.733 | 0.732 | 0.937 | 0.940 | 0.216 | 0.216 |
| | | M + 2 | 0.002 | 0.000 | 0.004 | 0.000 | 0.003 | 0.000 |
| Glx | [M-57]+ | M + 0 | 0.000 | 0.013 | 0.000 | 0.002 | 0.550 | 0.551 |
| | | M + 1 | 0.314 | 0.289 | 0.084 | 0.093 | 0.366 | 0.385 |
| | | M + 2 | 0.686 | 0.699 | 0.908 | 0.906 | 0.082 | 0.065 |
| | | M + 3 | 0.000 | 0.000 | 0.007 | 0.000 | 0.001 | 0.000 |
| | | M + 4 | 0.000 | 0.000 | 0.001 | 0.000 | 0.001 | 0.000 |
| | | M + 5 | 0.000 | 0.000 | 0.000 | 0.000 | 0.001 | 0.000 |
| | [M-85]+ | M + 0 | 0.045 | 0.047 | 0.020 | 0.022 | 0.706 | 0.706 |
| | | M + 1 | 0.953 | 0.953 | 0.970 | 0.978 | 0.291 | 0.294 |
| | | M + 2 | 0.001 | 0.000 | 0.009 | 0.000 | 0.001 | 0.000 |
| | | M + 3 | 0.001 | 0.000 | 0.001 | 0.000 | 0.001 | 0.000 |

| | | | | | | | | |
|---|---|---|---|---|---|---|---|---|
| | | M + 4 | 0.001 | 0.000 | 0.001 | 0.000 | 0.001 | 0.000 |
| | [M-159]$^+$ | M + 0 | 0.051 | 0.047 | 0.018 | 0.022 | 0.701 | 0.706 |
| | | M + 1 | 0.945 | 0.953 | 0.973 | 0.978 | 0.297 | 0.294 |
| | | M + 2 | 0.001 | 0.000 | 0.006 | 0.000 | 0.001 | 0.000 |
| | | M + 3 | 0.002 | 0.000 | 0.001 | 0.000 | 0.001 | 0.000 |
| | | M + 4 | 0.001 | 0.000 | 0.001 | 0.000 | 0.001 | 0.000 |
| Gly | [M-57]$^+$ | M + 0 | 0.264 | 0.268 | 0.059 | 0.060 | 0.907 | 0.906 |
| | | M + 1 | 0.735 | 0.732 | 0.938 | 0.940 | 0.092 | 0.094 |
| | | M + 2 | 0.001 | 0.000 | 0.003 | 0.000 | 0.001 | 0.000 |
| | [M-85]$^+$ | M + 0 | 0.999 | 1.000 | 0.994 | 1.000 | 0.999 | 1.000 |
| | | M + 1 | 0.001 | 0.000 | 0.006 | 0.000 | 0.001 | 0.000 |
| Ile | [M-85]$^+$ | M + 0 | 0.267 | 0.267 | 0.074 | 0.074 | 0.772 | 0.780 |
| | | M + 1 | 0.718 | 0.733 | 0.898 | 0.926 | 0.219 | 0.220 |
| | | M + 2 | 0.008 | 0.000 | 0.013 | 0.000 | 0.008 | 0.000 |
| | | M + 3 | 0.006 | 0.000 | 0.012 | 0.000 | 0.001 | 0.000 |
| | | M + 4 | 0.001 | 0.000 | 0.001 | 0.000 | 0.001 | 0.000 |
| | | M + 5 | 0.001 | 0.000 | 0.001 | 0.000 | 0.001 | 0.000 |
| | [M-159]$^+$ | M + 0 | 0.271 | 0.267 | 0.079 | 0.074 | 0.779 | 0.780 |
| | | M + 1 | 0.717 | 0.733 | 0.895 | 0.926 | 0.208 | 0.220 |
| | | M + 2 | 0.004 | 0.000 | 0.015 | 0.000 | 0.005 | 0.000 |
| | | M + 3 | 0.002 | 0.000 | 0.002 | 0.000 | 0.002 | 0.000 |
| | | M + 4 | 0.001 | 0.000 | 0.001 | 0.000 | 0.001 | 0.000 |
| | | M + 5 | 0.006 | 0.000 | 0.007 | 0.000 | 0.005 | 0.000 |
| Leu | [M-85]$^+$ | M + 0 | 0.994 | 1.000 | 0.979 | 1.000 | 0.993 | 1.000 |
| | | M + 1 | 0.001 | 0.000 | 0.014 | 0.000 | 0.004 | 0.000 |
| | | M + 2 | 0.004 | 0.000 | 0.004 | 0.000 | 0.001 | 0.000 |
| | | M + 3 | 0.001 | 0.000 | 0.001 | 0.000 | 0.001 | 0.000 |
| | | M + 4 | 0.000 | 0.000 | 0.001 | 0.000 | 0.001 | 0.000 |
| | | M + 5 | 0.000 | 0.000 | 0.001 | 0.000 | 0.001 | 0.000 |
| | [M-159]$^+$ | M + 0 | 0.989 | 1.000 | 0.973 | 1.000 | 0.986 | 1.000 |
| | | M + 1 | 0.000 | 0.000 | 0.015 | 0.000 | 0.002 | 0.000 |
| | | M + 2 | 0.004 | 0.000 | 0.004 | 0.000 | 0.007 | 0.000 |
| | | M + 3 | 0.005 | 0.000 | 0.005 | 0.000 | 0.003 | 0.000 |
| | | M + 4 | 0.001 | 0.000 | 0.001 | 0.000 | 0.001 | 0.000 |
| | | M + 5 | 0.001 | 0.000 | 0.001 | 0.000 | 0.001 | 0.000 |
| Phe | [M-57]$^+$ | M + 0 | 0.000 | 0.023 | 0.000 | 0.000 | 0.753 | 0.744 |

| | | | | | | | | |
|---|---|---|---|---|---|---|---|---|
| | | M + 1 | 0.191 | 0.177 | 0.017 | 0.015 | 0.213 | 0.231 |
| | | M + 2 | 0.450 | 0.439 | 0.177 | 0.193 | 0.029 | 0.024 |
| | | M + 3 | 0.358 | 0.360 | 0.790 | 0.792 | 0.002 | 0.001 |
| | | M + 4 | 0.000 | 0.000 | 0.015 | 0.000 | 0.001 | 0.000 |
| | | M + 5 | 0.000 | 0.000 | 0.000 | 0.000 | 0.001 | 0.000 |
| | | M + 6 | 0.000 | 0.000 | 0.000 | 0.000 | 0.000 | 0.000 |
| | | M + 7 | 0.000 | 0.000 | 0.000 | 0.000 | 0.001 | 0.000 |
| | | M + 8 | 0.000 | 0.000 | 0.000 | 0.000 | 0.000 | 0.000 |
| | | M + 9 | 0.000 | 0.000 | 0.000 | 0.000 | 0.000 | 0.000 |
| | $[M-85]^+$ | M + 0 | 0.063 | 0.088 | 0.000 | 0.006 | 0.808 | 0.821 |
| | | M + 1 | 0.430 | 0.420 | 0.138 | 0.151 | 0.171 | 0.170 |
| | | M + 2 | 0.496 | 0.492 | 0.827 | 0.842 | 0.016 | 0.009 |
| | | M + 3 | 0.006 | 0.000 | 0.020 | 0.000 | 0.001 | 0.000 |
| | | M + 4 | 0.003 | 0.000 | 0.013 | 0.000 | 0.001 | 0.000 |
| | | M + 5 | 0.001 | 0.000 | 0.001 | 0.000 | 0.001 | 0.000 |
| | | M + 6 | 0.000 | 0.000 | 0.000 | 0.000 | 0.001 | 0.000 |
| | | M + 7 | 0.001 | 0.000 | 0.000 | 0.000 | 0.001 | 0.000 |
| | | M + 8 | 0.000 | 0.000 | 0.000 | 0.000 | 0.001 | 0.000 |
| | $[M-159]^+$ | M + 0 | 0.067 | 0.088 | 0.001 | 0.006 | 0.787 | 0.821 |
| | | M + 1 | 0.425 | 0.420 | 0.143 | 0.151 | 0.159 | 0.170 |
| | | M + 2 | 0.489 | 0.492 | 0.825 | 0.842 | 0.019 | 0.009 |
| | | M + 3 | 0.005 | 0.000 | 0.019 | 0.000 | 0.007 | 0.000 |
| | | M + 4 | 0.005 | 0.000 | 0.005 | 0.000 | 0.005 | 0.000 |
| | | M + 5 | 0.003 | 0.000 | 0.002 | 0.000 | 0.005 | 0.000 |
| | | M + 6 | 0.002 | 0.000 | 0.002 | 0.000 | 0.005 | 0.000 |
| | | M + 7 | 0.002 | 0.000 | 0.001 | 0.000 | 0.006 | 0.000 |
| | | M + 8 | 0.003 | 0.000 | 0.001 | 0.000 | 0.007 | 0.000 |
| | $f302^+$ | M + 0 | 0.268 | 0.268 | 0.055 | 0.060 | 0.909 | 0.906 |
| | | M + 1 | 0.732 | 0.732 | 0.943 | 0.940 | 0.090 | 0.094 |
| | | M + 2 | 0.000 | 0.000 | 0.001 | 0.000 | 0.001 | 0.000 |
| Ser | $[M-57]^+$ | M + 0 | 0.259 | 0.268 | 0.052 | 0.060 | 0.904 | 0.906 |
| | | M + 1 | 0.740 | 0.732 | 0.940 | 0.940 | 0.095 | 0.094 |
| | | M + 2 | 0.001 | 0.000 | 0.006 | 0.000 | 0.001 | 0.000 |
| | | M + 3 | 0.001 | 0.000 | 0.001 | 0.000 | 0.001 | 0.000 |
| | $[M-85]^+$ | M + 0 | 1.000 | 1.000 | 0.989 | 1.000 | 0.993 | 1.000 |
| | | M + 1 | 0.000 | 0.000 | 0.010 | 0.000 | 0.006 | 0.000 |

| | | | | | | | | |
|---|---|---|---|---|---|---|---|---|
| | | M + 2 | 0.000 | 0.000 | 0.001 | 0.000 | 0.001 | 0.000 |
| | [M-159]$^+$ | M + 0 | 0.994 | 1.000 | 0.989 | 1.000 | 0.994 | 1.000 |
| | | M + 1 | 0.002 | 0.000 | 0.007 | 0.000 | 0.004 | 0.000 |
| | | M + 2 | 0.004 | 0.000 | 0.004 | 0.000 | 0.002 | 0.000 |
| | f302$^+$ | M + 0 | 0.268 | 0.268 | 0.065 | 0.060 | 0.909 | 0.906 |
| | | M + 1 | 0.732 | 0.732 | 0.935 | 0.940 | 0.087 | 0.094 |
| | | M + 2 | 0.000 | 0.000 | 0.000 | 0.000 | 0.004 | 0.000 |
| Val | [M-57]$^+$ | M + 0 | 0.268 | 0.268 | 0.064 | 0.060 | 0.992 | 0.992 |
| | | M + 1 | 0.730 | 0.732 | 0.924 | 0.940 | 0.006 | 0.008 |
| | | M + 2 | 0.001 | 0.000 | 0.010 | 0.000 | 0.001 | 0.000 |
| | | M + 3 | 0.000 | 0.000 | 0.001 | 0.000 | 0.001 | 0.000 |
| | | M + 4 | 0.000 | 0.000 | 0.001 | 0.000 | 0.001 | 0.000 |
| | | M + 5 | 0.001 | 0.000 | 0.001 | 0.000 | 0.001 | 0.000 |
| | [M-85]$^+$ | M + 0 | 0.980 | 1.000 | 0.961 | 1.000 | 0.973 | 1.000 |
| | | M + 1 | 0.006 | 0.000 | 0.016 | 0.000 | 0.024 | 0.000 |
| | | M + 2 | 0.013 | 0.000 | 0.021 | 0.000 | 0.001 | 0.000 |
| | | M + 3 | 0.001 | 0.000 | 0.001 | 0.000 | 0.001 | 0.000 |
| | | M + 4 | 0.000 | 0.000 | 0.001 | 0.000 | 0.001 | 0.000 |
| | [M-159]$^+$ | M + 0 | 0.970 | 1.000 | 0.950 | 1.000 | 0.962 | 1.000 |
| | | M + 1 | 0.001 | 0.000 | 0.014 | 0.000 | 0.002 | 0.000 |
| | | M + 2 | 0.004 | 0.000 | 0.004 | 0.000 | 0.004 | 0.000 |
| | | M + 3 | 0.023 | 0.000 | 0.028 | 0.000 | 0.030 | 0.000 |
| | | M + 4 | 0.003 | 0.000 | 0.004 | 0.000 | 0.002 | 0.000 |

**Table S4.** Best fitted fluxes and the 95% confidence intervals under acetate conditions.

| ID | Reaction | Best fit | | 95% confidence interval | |
|---|---|---|---|---|---|
| | | Net flux (mmol $g^{-1}$ $h^{-1}$) | Relative flux (%) | Lower limit (mmol $g^{-1}$ $h^{-1}$) | Upper limit (mmol $g^{-1}$ $h^{-1}$) |
| r1 | Subs_C2 --> AcCoA | 15.9 | 100.0 | 15.7 | 16.0 |
| r2 | AcCoA + OAA --> IsoCit | 10.2 | 64.3 | 10.0 | 10.4 |
| r3 | IsoCit --> AKG + CO2_in | 5.0 | 31.2 | 4.7 | 5.1 |
| r4 | AKG --> Sym_SUC + CO2_in | 4.5 | 28.4 | 4.3 | 4.7 |
| r5 | Sym_SUC <=> MAL | 9.8 | 61.5 | 9.5 | 9.9 |
| r6 | MAL <=> OAA | 11.2 | 70.3 | 10.0 | 13.4 |
| r7 | IsoCit + AcCoA --> MAL + Sym_SUC | 5.3 | 33.1 | 5.1 | 5.4 |
| r8 | PEP + CO2_in <=> OAA | 0.0 | 0.0 | -2.1 | 1.0 |
| r9 | MAL --> PYR + CO2_in | 3.9 | 24.3 | 1.8 | 5.0 |
| r10 | Subs_C3 --> PYR | 0.0 | 0.0 | 0.0 | 0.0 |
| r11 | PYR --> PEP | 2.1 | 12.9 | 0.0 | 3.0 |
| r12 | PEP <=> PGP | 1.7 | 11.0 | 1.7 | 1.7 |
| r13 | PGP <=> G3P | 1.1 | 7.0 | 1.1 | 1.1 |
| r14 | G3P <=> DHAP | 0.4 | 2.6 | 0.4 | 0.4 |
| r15 | DHAP + G3P <=> FBP | 0.4 | 2.6 | 0.4 | 0.4 |
| r16 | FBP --> F6P | 0.4 | 2.6 | 0.4 | 0.4 |
| r17 | F6P <=> G6P | 0.1 | 0.5 | 0.1 | 0.1 |
| r18 | PYR --> AcCoA + CO2_in | 0.6 | 3.9 | 0.5 | 0.8 |
| r19 | Ru5P <=> X5P | -0.3 | -1.9 | -0.3 | -0.3 |
| r20 | Ru5P <=> R5P | 0.3 | 1.9 | 0.3 | 0.3 |
| r21 | R5P + X5P <=> S7P + G3P | -0.1 | -0.5 | -0.1 | -0.1 |
| r22 | G3P + S7P <=> F6P + E4P | -0.1 | -0.5 | -0.1 | -0.1 |
| r23 | E4P + X5P <=> F6P + G3P | -0.2 | -1.4 | -0.2 | -0.2 |
| r24 | PGP --> Ser | 0.6 | 4.0 | 0.6 | 0.6 |
| r25 | Ser <=> Gly + THF_in | 0.2 | 1.5 | 0.2 | 0.2 |
| r26 | OAA --> ALA + CO2_in | 0.2 | 1.3 | 0.0 | 0.2 |
| r27 | PYR --> ALA | 0.0 | 0.0 | 0.0 | 0.2 |
| r28 | Subs_CO2 <=> CO2_in | -14.2 | -89.1 | -14.5 | -13.7 |
| r29 | Subs_THF <=> THF_in | -0.2 | -1.5 | -0.2 | -0.2 |
| r30 | Biomass | 0.4 | 2.6 | | |

**Table S5.** Best fitted fluxes and the 95% confidence intervals under ethanol conditions.

| ID | Reaction | Best fit | | 95% confidence interval | |
|---|---|---|---|---|---|
| | | Net flux (mmol $g^{-1}$ $h^{-1}$) | Relative flux (%) | Lower limit (mmol $g^{-1}$ $h^{-1}$) | Upper limit (mmol $g^{-1}$ $h^{-1}$) |
| r1 | Subs_C2 --> AcCoA | 14.0 | 100.0 | 13.5 | 14.1 |
| r2 | AcCoA + OAA --> IsoCit | 6.9 | 49.4 | 6.4 | 7.0 |
| r3 | IsoCit --> AKG + CO2_in | 1.0 | 6.8 | 0.6 | 1.1 |
| r4 | AKG --> Sym_SUC + CO2_in | 0.4 | 2.9 | 0.0 | 0.6 |
| r5 | Sym_SUC <=> MAL | 6.4 | 45.5 | 5.8 | 6.5 |
| r6 | MAL <=> OAA | 9.6 | 68.8 | -988.4 | 10.7 |
| r7 | IsoCit + AcCoA --> MAL + Sym_SUC | 6.0 | 42.6 | 5.8 | 6.1 |
| r8 | PEP + CO2_in <=> OAA | -1.8 | -12.7 | -2.6 | 996.0 |
| r9 | MAL --> PYR + CO2_in | 2.7 | 19.3 | 1.7 | 1000.0 |
| r10 | Subs_C3 --> PYR | 0.0 | 0.0 | 0.0 | 0.0 |
| r11 | PYR --> PEP | 0.8 | 5.6 | 0.0 | 998.5 |
| r12 | PEP <=> PGP | 2.2 | 15.6 | 2.2 | 2.2 |
| r13 | PGP <=> G3P | 1.4 | 10.0 | 1.4 | 1.4 |
| r14 | G3P <=> DHAP | 0.5 | 3.7 | 0.5 | 0.5 |
| r15 | DHAP + G3P <=> FBP | 0.5 | 3.7 | 0.5 | 0.5 |
| r16 | FBP --> F6P | 0.5 | 3.7 | 0.5 | 0.5 |
| r17 | F6P <=> G6P | 0.1 | 0.8 | 0.1 | 0.1 |
| r18 | PYR --> AcCoA + CO2_in | 0.2 | 1.2 | 0.0 | 0.3 |
| r19 | Ru5P <=> X5P | -0.4 | -2.7 | -0.4 | -0.4 |
| r20 | Ru5P <=> R5P | 0.4 | 2.7 | 0.4 | 0.4 |
| r21 | R5P + X5P <=> S7P + G3P | -0.1 | -0.7 | -0.1 | -0.1 |
| r22 | G3P + S7P <=> F6P + E4P | -0.1 | -0.7 | -0.1 | -0.1 |
| r23 | E4P + X5P <=> F6P + G3P | -0.3 | -2.0 | -0.3 | -0.3 |
| r24 | PGP --> Ser | 0.8 | 5.6 | 0.8 | 0.8 |
| r25 | Ser <=> Gly + THF_in | 0.3 | 2.2 | 0.3 | 0.3 |
| r26 | OAA --> ALA + CO2_in | 0.0 | 0.0 | 0.0 | 0.2 |
| r27 | PYR --> ALA | 0.3 | 1.8 | 0.1 | 0.3 |
| r28 | Subs_CO2 <=> CO2_in | -6.0 | -42.9 | -6.2 | -5.0 |
| r29 | Subs_THF <=> THF_in | -0.3 | -2.2 | -0.3 | -0.3 |
| r30 | Biomass | 0.5 | 3.8 | | |

**Table S6.** Best fitted fluxes and the 95% confidence intervals under ethanol and lactate conditions.

| ID | Reaction | Best fit | 95% confidence interval | |
|---|---|---|---|---|
| | | Net flux (mmol $g^{-1}$ $h^{-1}$) | Lower limit (mmol $g^{-1}$ $h^{-1}$) | Upper limit (mmol $g^{-1}$ $h^{-1}$) |
| r1 | Subs_C2 --> AcCoA | 14.0 | 13.5 | 14.1 |
| r2 | AcCoA + OAA --> IsoCit | 6.9 | 6.4 | 7.0 |
| r3 | IsoCit --> AKG + CO2_in | 1.0 | 0.6 | 1.1 |
| r4 | AKG --> Sym_SUC + CO2_in | 0.4 | 0.0 | 0.6 |
| r5 | Sym_SUC <=> MAL | 6.4 | 5.8 | 6.5 |
| r6 | MAL <=> OAA | 9.6 | -988.4 | 10.7 |
| r7 | IsoCit + AcCoA --> MAL + Sym_SUC | 6.0 | 5.8 | 6.1 |
| r8 | PEP + CO2_in <=> OAA | -1.8 | -2.6 | 996.0 |
| r9 | MAL --> PYR + CO2_in | 2.7 | 1.7 | 1000.0 |
| r10 | Subs_C3 --> PYR | 0.0 | 0.0 | 0.0 |
| r11 | PYR --> PEP | 0.8 | 0.0 | 998.5 |
| r12 | PEP <=> PGP | 2.2 | 2.2 | 2.2 |
| r13 | PGP <=> G3P | 1.4 | 1.4 | 1.4 |
| r14 | G3P <=> DHAP | 0.5 | 0.5 | 0.5 |
| r15 | DHAP + G3P <=> FBP | 0.5 | 0.5 | 0.5 |
| r16 | FBP --> F6P | 0.5 | 0.5 | 0.5 |
| r17 | F6P <=> G6P | 0.1 | 0.1 | 0.1 |
| r18 | PYR --> AcCoA + CO2_in | 0.2 | 0.0 | 0.3 |
| r19 | Ru5P <=> X5P | -0.4 | -0.4 | -0.4 |
| r20 | Ru5P <=> R5P | 0.4 | 0.4 | 0.4 |
| r21 | R5P + X5P <=> S7P + G3P | -0.1 | -0.1 | -0.1 |
| r22 | G3P + S7P <=> F6P + E4P | -0.1 | -0.1 | -0.1 |
| r23 | E4P + X5P <=> F6P + G3P | -0.3 | -0.3 | -0.3 |
| r24 | PGP --> Ser | 0.8 | 0.8 | 0.8 |
| r25 | Ser <=> Gly + THF_in | 0.3 | 0.3 | 0.3 |
| r26 | OAA --> ALA + CO2_in | 0.0 | 0.0 | 0.2 |
| r27 | PYR --> ALA | 0.3 | 0.1 | 0.3 |
| r28 | Subs_CO2 <=> CO2_in | -6.0 | -6.2 | -5.0 |
| r29 | Subs_THF <=> THF_in | -0.3 | -0.3 | -0.3 |
| r30 | Biomass | 0.5 | | |
